# As above, so below: Whole transcriptome profiling supports the continuum hypothesis of avian dorsal and ventral pallium organization

**DOI:** 10.1101/2020.11.13.375055

**Authors:** Gregory Gedman, Bettina Haase, Gillian Durieux, Matthew Biegler, Olivier Fedrigo, Erich D. Jarvis

## Abstract

Over the last two decades, beginning with the Avian Brain Nomenclature Forum in 2000, major revisions have been made to our understanding of the organization and nomenclature of the avian brain. However, there are still unresolved questions on avian pallial organization, particularly whether the cells above the ventricle represent different populations to those below it. Concerns included limited number of genes profiled, biased selection of genes, and potential independent origins of cell types in different parts of the brain. Here we test two competing hypotheses, using RNA sequencing to profile the transcriptomes of the major avian pallial subdivisions dorsal and ventral to the ventricle boundary, and a new zebra finch genome assembly containing about 22,000 annotated, complete genes. We found that the transcriptomes of neural populations below and above the ventricle were remarkably similar. What had been previously named hyperpallium densocellulare above the ventricle had nearly the same molecular profile as the mesopallium below it; the hyperpallium apicale above was highly similar to the nidopallium below; the primary sensory intercalated hyperpallium apicale above was most similar to the sensory population below, although more divergent than the other populations were to each other. These shared population expression profiles define unique functional specializations in anatomical structure development, synaptic transmission, signaling, and neurogenesis. These findings support the continuum hypothesis of avian brain subdivisions above and below the ventricle space, with the pallium as a whole consisting of four major cell populations instead of seven and has some profound implications for our understanding of vertebrate brain evolution.

## 1. INTRODUCTION

More than 120 years ago, some of the founders of comparative neurobiology proposed that the non-mammalian telencephalon consisted of mostly basal ganglia, homologous to the mammalian striatum and globus pallidus (Edinger, 1888; Edinger, 1908; Ariëns Kappers, 1922). This was the dominate view until the late 1960s, when the use of histochemical markers led to an alternative hypothesis that some of the striatal regions were really homologous to cell populations in the mammalian cortex (Karten, 1969). In the early 2000s, The Avian Brain Nomenclature Forum was formed, a consortium that evaluated the past century of findings and performed additional experiments, to develop a revised new nomenclature. They concluded that most of the striatum, the dorsal 2/3rds of the avian telencephalon was organized into distinct cell type subdivision broadly homologous to the developing mammalian pallium, inclusive of the 6-layered cortex, claustrum, and pallial amygdala (Reiner *et al*., 2004; Jarvis *et al*., 2005). As a result, the Forum, with support from the broader neuroscience community, developed a revised nomenclature that more accurately reflected homologies between the avian brain with those of mammals and potentially other vertebrates; this new nomenclature considered the pallial subdivisions on either side of the ventricular divide as uniquely different from each other (**Fig. 1a**).

**FIGURE 1.**
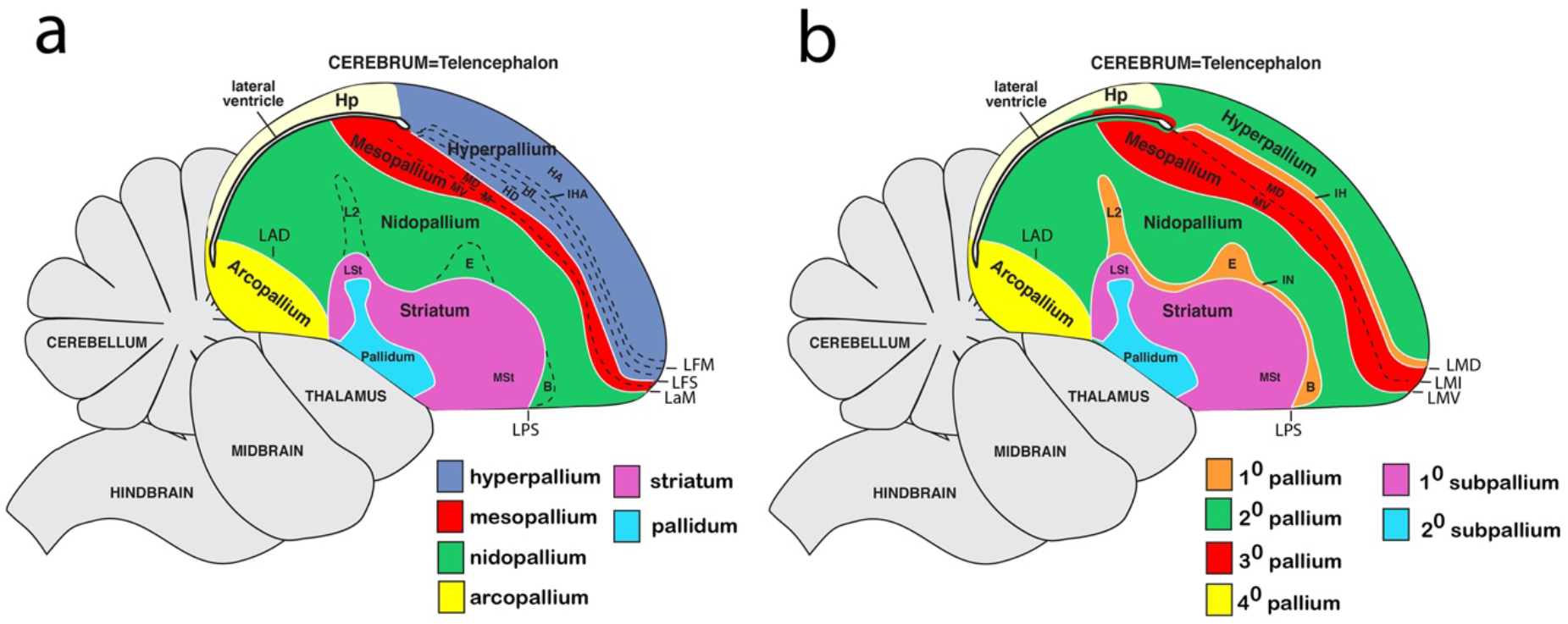
Two competing hypotheses on avian brain organization. (**a**) Dorsal and ventral pallium distinction hypothesis. (**b**) Dorsal and ventral pallium continuum hypothesis. Brain regions are colored according to similar cell populations according to each hypothesis. The lamina frontralis superior (LFS) in (white line in b) or lamina mesopallialus intermediate (LMI) (dashed line b) is the remaining vestigial lateral ventricle space that has become condensed in adults but is still connected with the more posterior lateral ventricle space shown below the hippocampus (Hp). Figure modified from Jarvis et al. (2013).

Despite this advance, a new model proposed nearly a decade later sparked new debate in our understanding of avian telencephalon organization. Jarvis et al. (2013) and Chen et al. (2013) proposed that what the Forum had revised to different hyperpallium populations above the lateral ventricle (**Fig. 1a, blue;** also called the Wulst), were in fact continuous with cell populations below the ventricle: the mesopallium, nidopallium, and sensory pallial areas of auditory L2, visual entopallium, and somatosensory basorostralis (**Fig. 1b, red, green, and orange;** also called the dorsal ventricular ridge, DVR). To explain this dorsal and ventral pallium continuum hypothesis, the findings suggests that as the ventricle space forms during embryonic development in birds, the cell types for these continuous subdivisions develop in tandem either above or below the ventricle, which then proliferate and simultaneously wrap around the ventricle, and later prior to hatching the ventricle space becomes partially occluded to form a separation between these continuous cell populations to form the dorsal and ventral pallium regions (Chen *et al*., 2013).

Although support for a dorsal and ventral mesopallium relationship above and below the ventricle has since been reported with RNA-Seq transcriptome data on embryonic chicken brains (Briscoe *et al*., 2018), the continuum hypothesis and associated “mirror image” view of avian brain organization above and below the ventricle has been met with some criticism. Despite the continuum model having been partly based on *in situ* hybridization expression profiles of over 50 genes, some have raised concern over the number and selection of the genes sampled (Montiel and Molnar 2013). There have also been concurrent gene expression studies that appeared to support the distinction hypothesis of cell populations above and below the ventricle (Dugas-Ford, Rowell and Ragsdale, 2012; Suzuki and Hirata, 2014; Belgard *et al*., 2013; Montiel and Molnár, 2013). One study found that the developmental expression profile of the *NR4A2* receptor only marked the mesopallium below the ventricle in birds, providing support to the distinction model (Puelles *et al*., 2016; Watson and Puelles, 2017). Another study proposed a hybrid hypothesis to try to reconcile the conflicting conclusions between these various studies (Wullimann, 2017). The ongoing debate over the precise organization of the avian brain makes it difficult to perform comparative and functional analyses within the avian brain, across vertebrate lineages, and between studies that rely different models (Lovell *et al*., 2020; Briscoe and Ragsdale, 2018; Puelles *et al*., 2016).

Here we attempt to resolve the debate by performing RNA-Seq transcriptome profiling on the main avian pallium populations in question to test the two main competing current hypotheses of avian brain organization (**Fig. 1**), using the zebra finch songbird (*Taeniopygia guttata*). The zebra finch belongs to the *Neoaves* clade, which makes up 95% of extant living bird species (Jarvis *et al*., 2014). Our comparative expression profiling of over 22,000 genes not only supports the continuum hypothesis of shared relationships of pallial populations below and above the lateral ventricle, it resolves discrepancies with the prior literature and reveals functional specializations specific to each population or combinations of populations (**Fig. 1b).**

## 1. MATERIALS AND METHODS

### 2.1 Animals subjects

Tissue samples were collected from four adult male zebra finches (~1-6 years old, **Table S1**). Animals were individually housed overnight in sound isolation chambers. Birds were euthanized in the dark two hours before the lights normally come on, by rapid decapitation within 1-2 min of handling, to limit activity-dependent gene expression changes in the brain (Wada *et al*., 2006; Feenders *et al*., 2008; Whitney *et al*., 2014). The brains were extracted, bisected sagittally, and each hemisphere frozen in Tissue-Tek OCT (Sakura, #4583) in a cryomold on dry ice. The entire procedure was performed within 5 minutes to reduce influence of activity-dependent genes and preserve RNA integrity.

### 2.2 Laser Capture Microscopy and RNA-Seq libraries/sequencing

One hemisphere/bird was sectioned on a cryostat at 12 μM and mounted on PEN membrane slides for Laser Capture Microscopy (LCM, Arcturus XT). In brief, one slide containing reference sections from each slide series was stained with Cresyl violet to aid in subdivision identification. PEN slides containing the sections of interest were individually dehydrated in serial alcohol baths from 50-100% and visualized under brightfield on the LCM (Arcturus Slide Prep Protocol #2). In addition to using the adjacent Cresyl violet stained section, axon bundles were visible in the LCM sections and used to help identify brain subdivisions. The region of interest was selected using a touch screen monitor and stylus pen and then laser dissected using the “Cut-and-Capture” method (Arcturus Instrument User Guide). First, a specialized cap with a microplastic film (Macro Caps: LCM0211) was placed on the tissue, then an infrared laser was used to melt the microplastic film to “capture” the tissue on the cap, and finally an ultraviolet laser was used to “cut” the region of interest from the larger tissue section. All tissue samples were dissected in under 30 minutes/slide to ensure high RNA integrity. A minimum RIN of 6 (mean = 7.1) was required for further processing for sequencing. RNA was isolated from each sample using the Picopure RNA Isolation kit (Ref: KIT0204), and stored at −80° until all samples were collected. Samples were randomized across batches (n=4) to minimize batch effects. However, some samples, specifically the intercalated nidopallium and hyperpallium, were collected several years apart in sections from the same animals, so additional analyses were performed to test and normalize for any potential batch effects. Samples were randomized into batches and cDNA was generated using the SMART-Seq Ultra Low-Input RNA kit for sequencing (Clonetech, Ref: 634891). Each sample library was prepped using the NEBNext Ultra II DNA Library Prep Kit (Cat: E7645L) and duel-indexed for sequencing using the NEBNext Multiplex Oligos for Illumina Set 2 (Cat: E6442S). RNA Sequencing of pair-end, 150bp reads was conducted on the NextSeq 500 system from Illumina.

### 2.3 Sequencing data quality control

Quality of all raw sequence reads were verified using FastQC (v0.11.5), trimming off low-quality (<QV30) and adapter sequences using fastq-mcf (v1.05). Reads were mapped to the newly assembled and annotated high-quality, long-read based, Vertebrate Project Genome (VGP) zebra finch genome (bTaeGut1_v1, RefSeq Accession: GCF_003957565.1). Transcript levels for each gene in each brain region were determined using Salmon (v0.14.1). A final gene x sample matrix was used as input for all downstream analyses. Housekeeping genes were empirically determined based on their expression variation, with any gene with a coefficient of variance (CV) of zero across all samples taken as housekeeping. The scater package in R (v1.12.2) was used to explore the effects of unwanted variations from known sources. Any significant sources of unwanted variation, like individual bird effect, were accounted for in all downstream analyses either by inclusion as a term in linear models or direct correction using the “removeBatchEffect” function from limma (v3.40.6).

### 2.4 Principle component, differential expression, and molecular anatomical cluster analyses

The raw gene x sample expression matrix was supplied to DESeq2 (v1.24.0) for differential expression testing between brain subdivisions. Following variance stabilization transformation, we performed principle component analysis (PCA) and plotted the first two components explaining a majority of the variance. For differential expression testing, a linear model was constructed to model brain subdivision and individual animals, and then each subdivision was contrasted to all others in a pairwise manner. Genes were considered differentially expressed if they passed multiple test corrections (q < 0.05). A dissimilarity matrix was generated from the total number of differentially expressed genes (i.e. degree of difference) for each comparison and results clustered using the “hclust” (method = “average”) function from R. A union set of all differentially expressed genes (DEGs) was taken; a similar clustering procedure was performed on the normalized counts to determined degree of shared expression between all samples for the genes with the strongest biological signal. Bootstrap resampling was also performed using 1,000 iterations of all DEGs using the pvclust package in R (v2.2-0). Importantly, there are frequent improvements to the zebra finch assembly/annotation, so currently uncharacterized genes (LOC IDs) may be annotated with gene symbols following the publication of this manuscript. All genes from this analysis (Table S2) can be searched in NCBI’s Genome Data Viewer (https://www.ncbi.nlm.nih.gov/genome/gdv/) under the zebra finch genome version used in this study (bTaeGut1_v1, RefSeq Accession: GCF_003957565.1) and the most up-to-date records will be displayed.

### 2.5 Weighted gene co-expression network construction

Gene networks were assessed following the tutorial for weighted gene network co-expression analysis using the WGCNA package (v1.69) in R. In brief, samples were screened using the “goodSamplegenes” function, resulting in 20,822 stably expressed genes identified across all samples. The weighted adjacency matrix was constructed using a soft-power threshold = 6. This adjacency matrix was converted into a topological overlap matrix (TOM), and gene modules were clustered by TOM-based dissimilarity (1-TOM). The minimum module size was set to 100 genes in order to reduce obtaining single sample modules specific to a bird.

### 2.6 Identification of brain subdivision gene co-expression modules

Module eigengenes (MEs) were calculated for each module, allowing the expression patterns of all genes in a module to be summarized to a single statistic. Each subdivision was coded as either a unique brain region or combination of regions depending on hypothesis, and significant modules were determined by correlation with each module’s eigengene vector using the “corPvalueStudent” function from the WGNCA package (Langfelder and Horvath, 2008). P-values were corrected for multiple tests using the “p.adj” function in R. Modules were considered highly significantly correlated for each brain subdivision if they exhibited an r^2^ value > 0.90 and a q value < 0.001. Gene ontology analysis was conducted to assess functional enrichment of each significant module with a *H. sapiens* background and custom expression background (mean expression < 10/all samples) using gProfileR2 (v0.1.9).

### 2.7 Hub gene identification for subdivision modules

Two metrics were used to determine significant hub genes for each brain subdivision module. First, gene significance (GS, strength of correlation to a region) was defined as the Pearson correlation of each gene expression value to each brain subdivision. P-values were calculated for each correlation and corrected for multiple tests as described above. Next, module membership (MM, strength of connectivity in the module) was determined using the Pearson correlation of each gene’s expression value with the eigengene vector for each module. Hub genes were defined by their significant correlation with a brain subdivision (absolute value of GS > 0.8) and strong connectivity with other module genes (absolute value of MM > 0.8). Networks of the top 50 interconnected hub genes were visualized using VizANT (v1.0). Uncharacterized genes (LOC IDs) were replaced with other aliases whenever possible using the rentrez package (v1.2.2).

## 2. RESULTS

### 3.1 Collecting basal transcriptome levels

Our goal was to determine the relationship between various subdivisions of the avian telencephalon according to two leading hypotheses of brain organization (**Fig. 1**) by measuring their transcriptomes at baseline. In order to compare different brain regions across different individuals, we needed to collect brain tissue from animals under carefully controlled conditions in order limit any confounding variables. Since up to 10% of the transcribed genome can be regulated in an activity-dependent manner, with different cell populations having different sets of regulated genes in different brain regions controlled by different behaviors (Jarvis *et al*., 2013; Whitney *et al*., 2014), we designed our experiment to reduce these confounding variables. Male zebra finches were kept alone in sound isolation chambers overnight to keep gene expression levels at steady state and only males were used to prevent any differences in sex chromosome expressed genes to confound our analyses. Sagittal sections were processed in order to reduce section as a variable, as more brain subdivisions are captured together in the sagittal plane relative to the more commonly cut coronal plane.

A series of sections were stained with Cresyl violet to help identify brain subdivisions. Adjacent sections were processed for laser capture microdissection (LCM), using brightfield microscopy to help further identity brain subdivision boundaries, and aid in avoiding accidentally contaminating samples with adjacent brain subdivisions (**Fig. 2**). In order to consider both hypotheses (**Fig. 1a,b**), we captured nine regions from dorsal and ventral pallial subdivisions. From the dorsal pallium regions (e.g. Wulst) we captured: 1) a visual region of the hyperpallium apicale (HA; aka hyperpallium, H; **Fig. 2a**); 2) a visual region of the underlying intercalated hyperpallium apicale (IHA; aka intercalated hyperpallium, IH; **Fig. 2b**); and 3) a visual region of the hyperpallium densocellure (HD; aka dorsal mesopallium, MD; **Fig. 2a**). From the ventral pallium regions (e.g. the_ dorsal ventricular ridge; DVR), we captured: 4) a motor region of the mesopallium (M; aka ventral mesopallium, MV; **Fig. 2a**); 5,6) anterior motor and posterior lateral auditory-motor regions of the nidopallium (AN and PLN, respectively; **Fig. 2c,d**); and 7) the Field L2 auditory portion of the intercalated nidopallium (IN; aka L2; **Fig. 2b**). We isolated two regions of the nidopallium to test for diversity within an accepted brain subdivision. We also isolated two regions from two other unique subdivisions accepted by both hypotheses: 8) a motor portion of the lateral intermediate arcopallium (LAI; **Fig. 2e**) and 9) a motor portion of the ventral striatum (VSt; **Fig. 2f**). A list of region abbreviations in the context of both hypotheses is provided in Figure 2. The motor and sensory functional designations of each subdivision were according to primary and reviewed findings in previous studies (Feenders *et al*., 2008; Jarvis *et al*., 2013). We could not find a fourth hyperpallial region between the formally named hyperpallium densocellare and intercalated hyperpallium apicale (**Figs. 1a, 2**), consistent with our previous findings in zebra finches and other avian species (Jarvis *et al*., 2013). The dissected motor regions of the nidopallium, arcopallium, and striatum were adjacent to the song nuclei of those brain subdivisions (**Fig. 2c,d,f**). After LCM dissection, total RNA was isolated from each sample in a randomized batch design. In cases where the minimum RNA integrity (RIN) value of 6 was not reached, then the procedure was repeated on another adjacent section.

**FIGURE 2.**
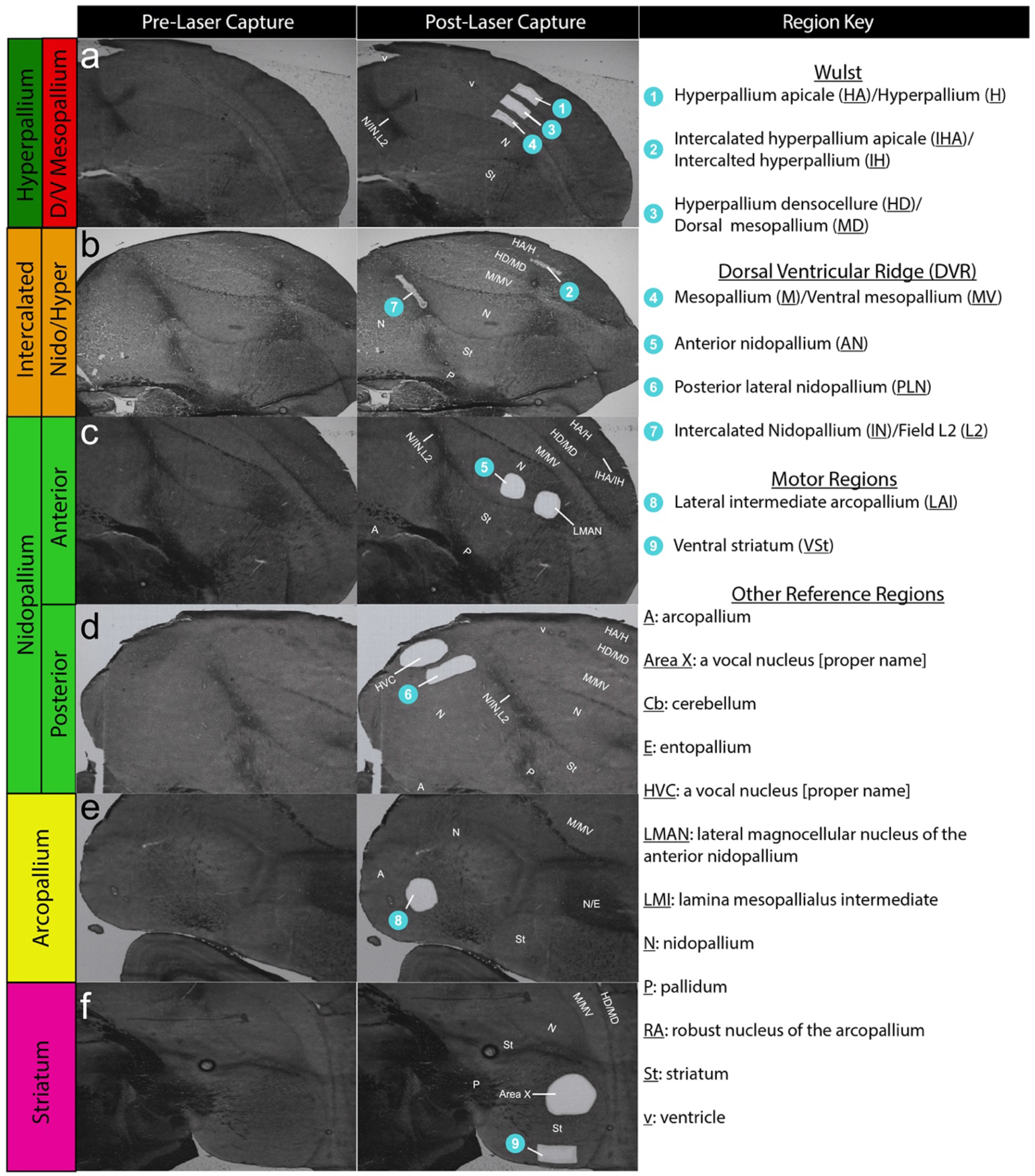
Example images of LCM dissections. (**a-e**) Brightfield sections showing the brain regions profiled before (left column) and after (center column) LCM dissections. Dissected regions are numbered (blue circles). Abbreviations for all relevant brain regions according to the distinction (**Fig. 1a**) and continuum hypotheses (**Fig. 1b**) are provided (right column). Some sections have additional dissections of song nuclei as part of another study in progress. Darker brain regions are due to increased myelination, some of which separate brain subdivisions via axon tracts.

RNA-Seq expression profiling was performed on these samples, with a sequencing depth of ~ 20 million reads per sample. These reads were aligned to a new more complete and error-free 2019 zebra finch genome assembly from the Vertebrate Genomes Project (VGP), containing 22,186 annotated genes, of which 17,438 are protein coding (Rhie *et al*., 2020). We were able to map nearly 98% of reads to transcripts from this new VGP assembly, maximizing the power of our analysis, compared to 87% to the old Sanger-based assembly (Warren *et al*., 2010; Rhie *et al*., 2020). Importantly, 91% of the reads mapped to unique loci (21,617 genes, > than one read in a sample) in the 2019 assembly (~84% in 2010 assembly), which were used for all downstream analyses.

### 3.2 Quality control and impact of bird specific transcriptome patterns

We first tested whether there were any batch affects or other technical influences on the gene expression patterns across all samples. Although the intercalated pallium samples were collected separately from the other samples in the study, we did not detect any major batch effects using a variety of approaches. For example, we assessed the distribution of the normalized expression counts for the intercalated samples relative to the rest of the samples and compared their quantiles in a Q-Q plot (**Fig. 3a**). We found that their distributions were nearly identical, indicating there was no systematic shift between collection groups. A plot of the relative log expression of all genes in each sample indicated no evidence of a systematic shift in global expression in any sample or collection group (**Fig. 3b**). We also assessed the distribution of house-keeping genes across both collection groups. We calculated the coefficient of variance (CV) for all batches in the first collection group, and empirically defined house-keeping genes as those with a CV = 0. About 81% of these genes showed stable expression in the second collection group, further supporting their consistency following normalization regardless of batch (**Fig. 3c**). Results presented further below suggest that the 19% difference in the housekeeping genes is biologically driven.

**FIGURE 3.**
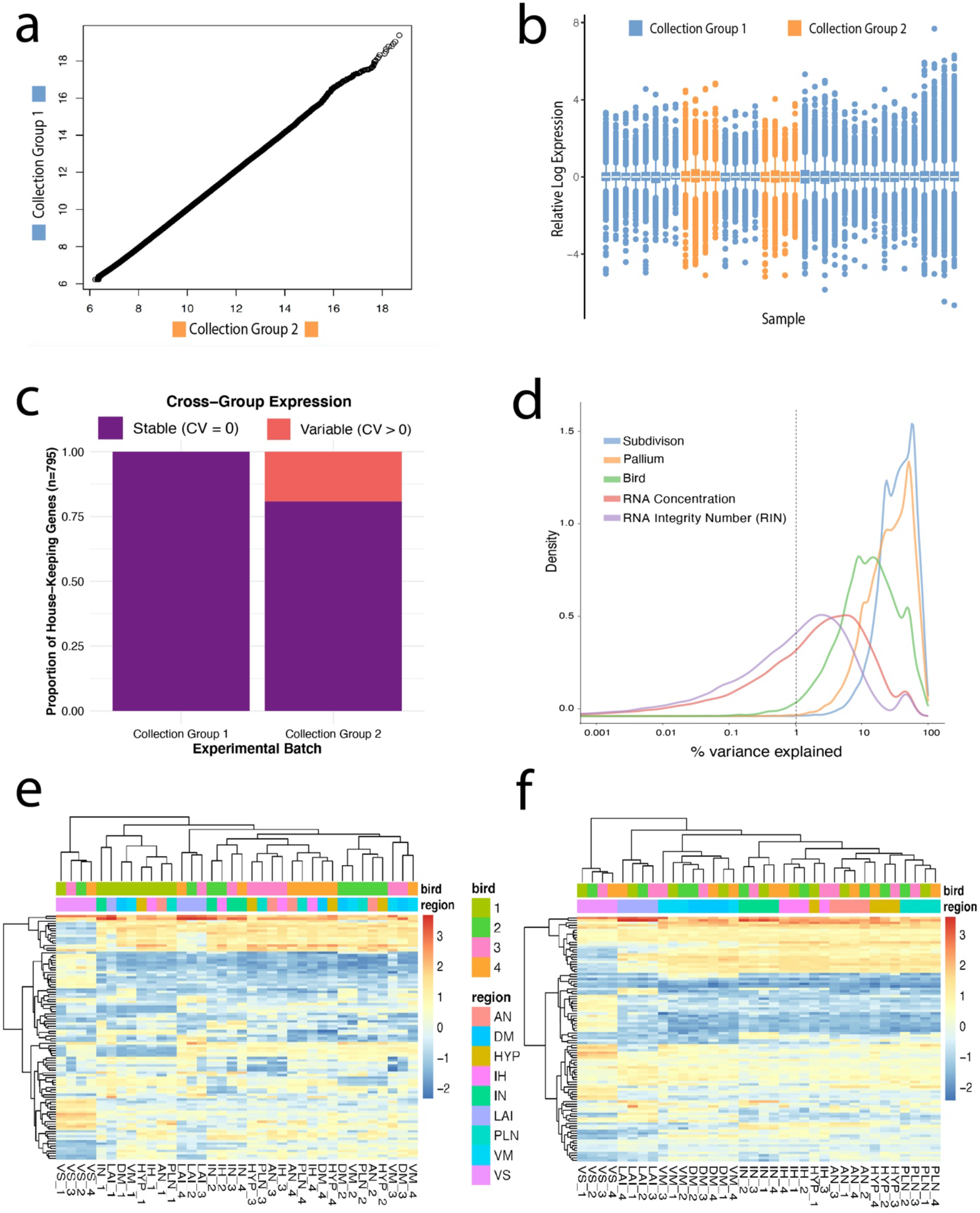
Minimal batch effects detected across collection groups, while individual bird is a strong source of unwanted variation. (**a**) Q-Q plot of normalized expression from collection group 1 (all non-intercalated samples) vs. collection group 2 (intercalated samples). The distributions look nearly identical. (**b**) Relative log expression plot of all samples colored by collection group. There is no evidence of a systematic shift in global expression in any sample. (**c**) Proportion of stable housekeeping genes (CV = 0) across the collection groups. (**d**) Density plot of variance explained by individual variables. Note brain subdivisions (blue line) and broader pallium subdivisions (orange line) together explain most of the variance seen in the data, individual bird (green line) accounts for a non-trivial amount of variance, and RNA concentration and RIN have little effect. (**e**) Heatmap of normalized expression of top 100 most variable genes. Regions cluster mostly by bird (top color bar) rather than brain subdivisions regions (bottom color bar). (**f**) After accounting for the bird effect as a covariate, these same genes exhibit robust clustering by brain region (bottom color bar).

We maximized the variance explained by our biological variables of interest (subdivision and pallium) by modeling and removing sources of unwanted variation. We found that brain subdivision, including broader pallium subdivisions, explained the vast majority (>90%) of the variance distribution (**Fig. 3d**). However, there was a strong effect peaking around 10% of variance from individual birds, and a weaker one with ~5% of variance associated with RNA concentration and quality (**Fig. 3d**). Despite the strong explanation by brain subdivision, hierarchical clustering of expression levels of the top 100 most variable genes in the data clustered samples more by individual bird than by brain region (**Fig. 3e**). Removing this individual effect from the data utilizing the “removeBatchEffect” function from R’s limma package resulted in robust clustering by brain region (**Fig. 3f**), highlighting the importance of controlling for individual animal variation before conducting downstream analyses. This bird effect was accounted for in all analyses in this study, either by direct removal from the normalized expression matrix (principle component analysis, gene network analysis) or by inclusion in the linear model for differential expression testing.

### 3.3 Shared brain molecular profiles between Wulst and DVR brain regions

Principle component analysis (PCA) of all 21,617 unique genes in the bird-normalized expression matrix from all samples revealed consistent clustering among most samples from the same brain region (**Fig. 4a**). The first principle component (PC) was explained by large differences between the striatal and pallial regions for all birds; the second PC was explained by differences among pallial regions, with the arcopallium being the most distinct, followed by the intercalated sensory regions (intercalated nidopallium and intercalated hyperpallium), followed by the other pallial regions (dorsal mesopallium, ventral mesopallium, hyperpallium, and nidopallium), which all grouped closer to each other (using the terminology of the continuum hypothesis). To further explore the clustering of the pallial populations in the Wulst and DVR surrounding the ventricle, we ran an additional PCA of pallial samples without the arcopallium and striatum. The Wulst samples above the ventricle (hyperpallium, intercalated hyperpallium, and dorsal mesopallium) did not form distinct clusters from the DVR samples below the ventricle (nidopallium, intercalated nidopallium, and ventral mesopallium; **Fig. 4b**). Instead, the samples from each subdivision in the Wulst clustered with one subdivision each in the DVR: hyperpallium with nidopallium; intercalated hyperpallium with intercalated nidopallium; and dorsal mesopallium with ventral mesopallium (**Fig. 4b**). Given these results, from here on we will primarily refer to the profiled brain regions using their names under the continuum hypothesis.

**FIGURE 4.**
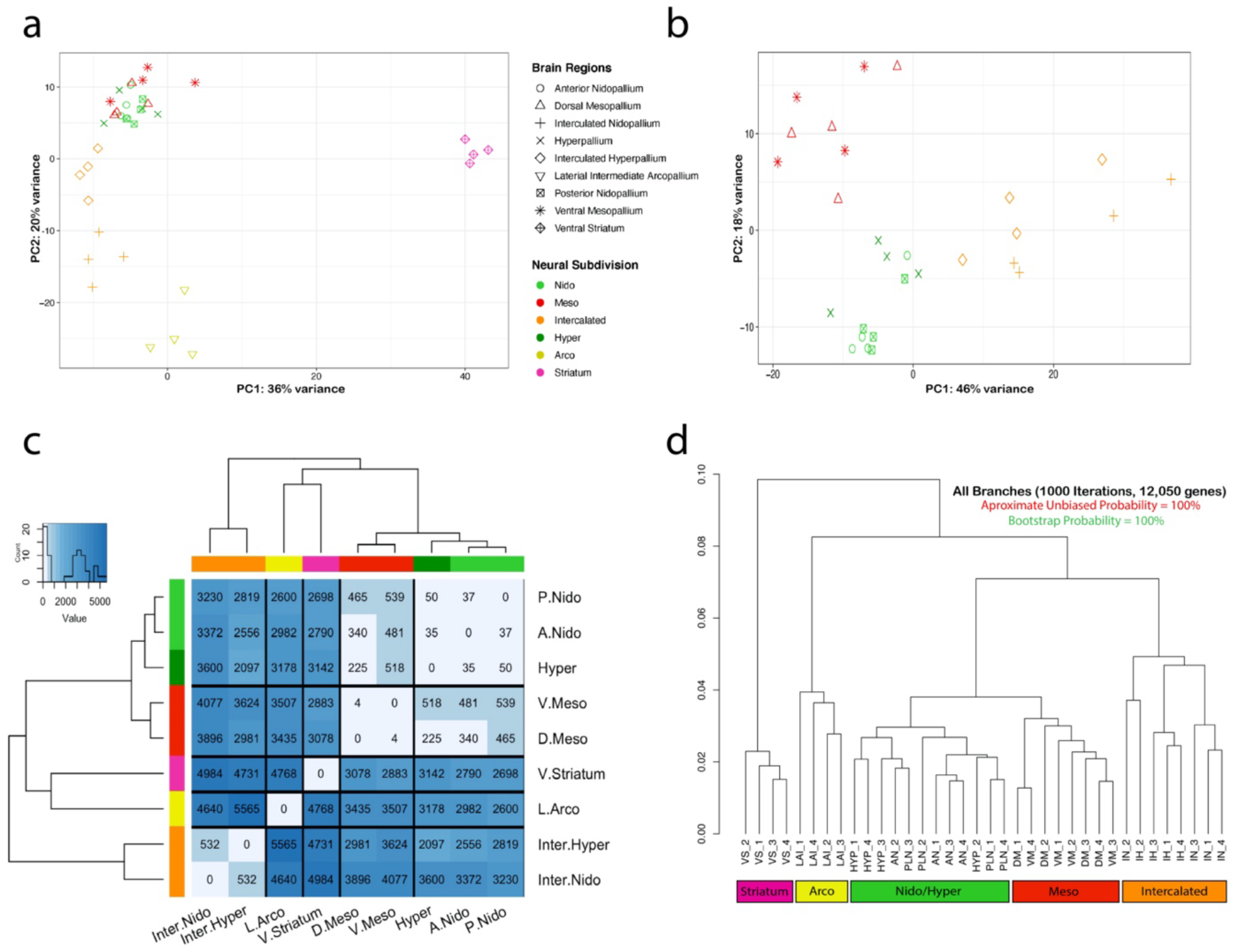
Molecular relationships between brain subdivisions. (**a**) PCA of whole transcriptome expression for all brain regions profiled. (**b**) PCA of whole transcriptome expression in nidopallium, hyperpallium, dorsal mesopallium, ventral mesopallium and intercalated regions. PCAs of whole transcriptome gene expression sample n=21,617 genes. Each point represents measurements from one bird. The symbol legend treats the brain regions above and below the ventricle as different in the context of the distinction hypothesis. The color treats them as similar in the context of the continuum hypothesis. Note that samples above the ventricle are not distinct but cluster with samples from below the ventricle. (**c**) Dissimilarity heatmap with cluster dendrograms of all differentially expressed genes in pairwise analyses of all brain subdivisions profiled. Heatmap is colored according to number of genes that are significantly differentially expressed at a false discovery rate (FDR) < 0.05. As samples cluster together according to the continuum hypothesis, the leaves of the tree are colored and labeled according to the continuum hypothesis. (**d**) Hierarchical clustering with approximate unbiased (au, red) and bootstrap probability (bp, green) p-values for all differentially expressed genes (n=12,050) across all samples. All approximate unbiased (au, red) and bootstrap values were 100% for all branches. See Figure 2 for brain region abbreviation list; numbers next to the abbreviations are individual birds.

In order assess similarities in the molecular specializations of each brain region, pairwise differential expression analysis was performed on all measured genes (n=21,617) for each subdivision and results were hierarchically clustered based on the total number of genes with significant differences. We first noted that two regions from the same brain subdivision (anterior and posterior nidopallium) differed by only 37 genes (0.1% of genes tested; **Fig. 4c**), setting a threshold of when to consider if two regions belonging to the same brain subdivision. Given this threshold, we looked at the regions above and below the ventricle (**Fig. 4c**). Specifically, we found that the dorsal mesopallium (aka hyperpallium densocellulare in **Fig. 1a**) above the ventricle was the most similar to the ventral mesopallium (aka mesopallium) below it, differing by only 4 genes, about 0.01% of the genes tested for differential expression (FDR < 0.05). The hyperpallium (aka hyperpallium apicale) above the ventricle was most similar to the anterior and posterior nidopallium below the ventricle, differing by only 35 and 50 genes, respectively (0.1% and 0.2% of the genes tested; **Fig. 4c**). Remarkably, the anterior nidopallium was more similar to the hyperpallium than it was to the posterior nidopallium, suggesting that two areas within one brain subdivision below the ventricle (nidopallium) are as different from each other as one of them is to a subdivision above the ventricle (hyperpallium). The intercalated hyperpallium (aka intercalated hyperpallium apicale in **Fig. 1a)** above the ventricle was the most similar to the intercalated nidopallium (Field L2) below the ventricle, but differing by many more genes (n = 532, 2.6% of genes tested; **Fig. 4c**). Despite this greater diversity, by comparison the intercalated hyperpallium and intercalated nidopallium are 4 and 6 times more different than the brain subdivisions they have long been assumed to belong, the hyperpallium (n = 2,097 genes; 10.4%) and nidopallium (n = 3,230 genes; 16.1%; **Fig. 4c**) respectively. These differences between the intercalated pallium regions with the nidopallium and hyperpallium are in the range of the number of genes that differ between well-established brain subdivisions (~3000-6000 genes; ~15-30%), including between pallial and striatal regions (**Fig. 4c**).

While this clustering approach details extent of total gene differences, it doesn’t reflect shared character of specializations, i.e. regions that have the same genes specialized in the same/opposite directions. In order to visualize brain region relationships for the genes with the most biological signal, we took the union set of all statistically significant, differentially expressed genes (n=12,050) and performed hierarchical clustering of the expression values with bootstrap sampling of all genes in all brain regions, keeping each sample of each bird independent (**Fig. 4d**). Remarkably, this phylo-gene expression tree showed a similar topology as that from using the 50 genes sampled in Jarvis et al. (2013). The striatum clustered away from the other pallium samples, with the arcopallium being the most distinct of the pallial regions. The remaining pallial samples clustered together in a pattern that supports the hypothesized continuous relationships, with 100% bootstrap probability support in all branches. Further, the nidopallium/hyperpallium and mesopallium regions formed a super cluster, revealing higher order relationships. In all three types of analyses (**Fig. 4**), we did not observe any clustering pattern that supported grouping the hyperpallium subdivisions above the ventricle (**Fig. 1a**) as more similar to each other. Rather, the gene expression clustering via PCA (**Fig. 4a,b**), differential gene expression (**Fig. 4c**), and phylogenetic bootstrapping similarities (**Fig. 4d**) were antithetical to the distinction hypothesis.

### 3.4 Validations by *in situ* hybridization

To validate our RNA-Seq findings and determine if the expression profiles we discovered are characteristic of the brain subdivisions, we analyzed available *in situ* hybridization profiles of 64 genes from various studies (Wada *et al*., 2004; Kubikova, Wada and Jarvis, 2010; Jarvis *et al*., 2013; Chen *et al*., 2013; Pfenning *et al*., 2014; Whitney *et al*., 2014) and the zebra finch expression atlas (Lovell *et al*., 2020) which had clear expression profiles and high-quality data (**Table S2**). This included searching for available *in situs* of all genes with significant differential RNA-Seq expression in the Wulst and DVR populations above and below the ventricle. We scored each of these gene’s patterns in pairwise differential expression results as True Positive (TP), True Negative (TN), False Positive (FP), or False Negative (FN), and calculated Accuracy = (TP+TN)/(TP+TN+FP+FN). These findings show that RNA-Seq accuracy was very high and is in concordance with known markers for these brain subdivisions (**Table 1**). The accuracy of the differential expression to known markers for the two mesopallium regions was 100% (**Table 1**). The accuracy between the hyperpallium and anterior nidopallium expression was ~97%, further supporting the similarity observed between these regions. The accuracy between the intercalated pallium and the posterior nidopallium and hyperpallium regions in which they reside, respectively was ~86% and 90%, while the accuracy between the two intercalated regions was 95%. By comparison the accuracy of expression between the well-established arcopallium versus striatum subdivisions was 89%. These findings indicate that the RNA-Seq gene expression comparisons between more similar brain subdivisions have higher accuracy according to our *insitu* hybridization analyses.

**TABLE 1:** Concordance of differential expression analyses with known control genes. True positive, false positive, true negative, and false negative values and the percent accuracy from those values are listed for 64 genes, between 10 brain subdivision comparisons that test the distinction versus continuum hypotheses their relationships. All RNA-Seq and *in situ* hybridization comparisons examined had a percent accuracy between 84-100%, adding confidence to the differential gene expression and hierarchical clustering results (**Fig. 4**). A: arcopallium, AN: anterior nidopallium, H: hyperpallium, IH: intercalated hyperpallium, IN: intercalated nidopallium, MD: dorsal mesopallium, MV: ventral mesopallium, PLN: posterior lateral nidopallium, St: Striatum.

| Region 1 | Region 2 | True Positive | False Positive | True Negative | False Negative | Percent Accuracy |
| --- | --- | --- | --- | --- | --- | --- |
| MD | MV | 1 | 0 | 63 | 0 | 100.0 |
| H | AN | 5 | 1 | 57 | 1 | 96.9 |
| AN | PLN | 3 | 0 | 59 | 2 | 96.9 |
| A | PLN | 34 | 2 | 28 | 0 | 96.9 |
| IH | IN | 10 | 2 | 51 | 1 | 95.3 |
| IN | PLN | 27 | 4 | 31 | 2 | 90.6 |
| PLN | MV | 20 | 1 | 37 | 6 | 89.1 |
| A | St | 45 | 6 | 12 | 1 | 89.1 |
| IH | H | 11 | 4 | 44 | 5 | 85.9 |
| H | MD | 14 | 1 | 40 | 9 | 84.4 |

Example validated genes of shared specialized expression between the Wulst and DVR regions included *SATB2* (Special AT-Rich Sequence-Binding Protein 2) and *CHRNA3* (Cholinergic Receptor Nicotinic Alpha 3 Subunit), both upregulated equally in the dorsal mesopallium and ventral mesopallium relative to other pallial regions (**Fig. 5a,b**). *KCTD12* (Potassium Channel Tetramerization Domain Containing 12) was equally upregulated in the hyperpallium and nidopallium, and not in the mesopallium regions (**Fig. 5c**). *SLC4A4* (Solute Carrier Family 4 Member 4), involved in the regulation of bicarbonate secretion and absorption, and intracellular pH, was confirmed with upregulation specific to the intercalated pallial regions (**Fig. 5d**).

**FIGURE 5.**
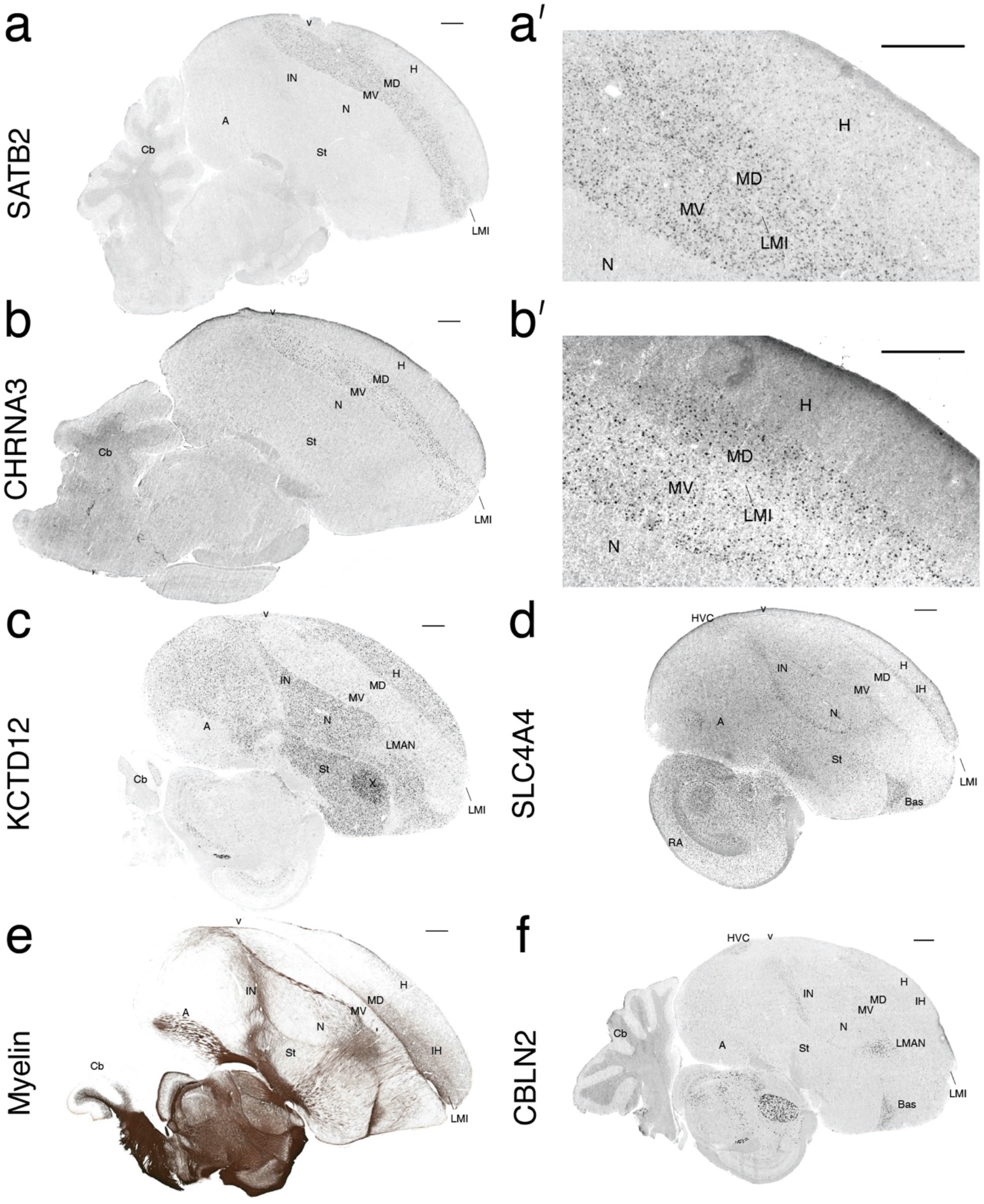
*In situ* hybridizations and myelin staining confirm RNA-Seq profiles and reveal full anatomical expression patterns. (**a**) *SATB2*, a mesopallium upregulated gene; **(a’**) higher power image shows that it also has sparse hyperpallium expression, not seen in the nidopallium. (**b**) *CHRNA3*, a mesopallium upregulated gene; **(b’)** higher power image shows that it does not have sparse expression in the hyperpallium. (**c**) *KCTD12*, a nidopallium and hyperpallium upregulated gene. (**d**) *SLC4A4*, an intercalated pallium upregulated gene. (**e**) Myelin stain correlating with increased expression of *MBP* in the anterior dorsal mesopallium (MD) relative to the ventral mesopallium (MV). (**f**) *CBLN2* is upregulated in the intercalated nidopallium but not in the intercalated hyperpallium. All *in situ* hybridization images are from the zebra finch gene expression atlas (Lovell *et al*., 2020) and downloaded as of August 2020. The myelin image is from the digital atlas of (Karten *et al*., 2013). All scale bars are 1mm.

Of the few rare genes that exhibited differential expression between the shared Wulst and DVR regions, we validated several. Using a myelination stain, we observed that myelin, whose major component is Myelin Basic Protein (MBP), was notably higher in the dorsal mesopallium relative to the ventral mesopallium, consistent with the RNA-Seq expression data (**Fig. 5e, Table S2**). The reason for this difference appeared to be a higher number of myelinated fibers coursing medial-laterally in the anterior half of the dorsal mesopallium. Notably, one of the other three genes with increased expression in dorsal mesopallium relative to ventral mesopallium, *NINJ2* (Ninjurin 2), has been shown to be differentially expressed in adult myelinating oligodendrocytes in comparative high-throughput microarray screens of mouse cortical cell types (Noroozi *et al*., 2019), further suggesting the principle difference in the mesopallium regions surrounding the ventricle is myelin based. *SATB2* was also one of the top five of the 36 genes with differential expression between the nidopallium and hyperpallium, and the gene with the highest expression difference in the hyperpallium relative to the nidopallium (**Table S2**). The *in situ* hybridization revealed that the reason for this difference was higher expression in sparsely labelled cells throughout the hyperpallium not found in the nidopallium (**Fig. 5a’**). This is similar to the pattern observed previously (Jarvis *et al*., 2013) with *SCUBE1* (Signal Peptide, CUB Domain, EGF-Like Domain Containing 1), which apparently was not strong enough to rise to the level of significance in the RNA-Seq data after multiple test corrections. We have also noticed a sparse hyperpallium expression pattern with *NR4A2* (Nuclear Receptor Subfamily 4 Group A Member 2) during activity-dependent induction in our companion study (Biegler et al., 2020 submitted). These findings hint at a sparse cell type unique to the hyperpallium relative to the nidopallium, which may come about through the migration of the nidopallium through the mesopallium during development (Chen *et al*., 2013). However, most mesopallium markers did not show similarly labelled sparse cells in the hyperpallium at baseline (**Fig. 5b’**). The top ranked gene with higher expression in the nidopallium relative to the hyperpallium was *NR2F2* (Nuclear Receptor Subfamily 2 Group F Member 2, a.k.a. *COUPTFII*), which had been previously identified as a nidopallium marker relative to the hyperpallium (Jarvis *et al*., 2013). The *CBLN2* (Cerebellin 2 Precursor) *in situ* pattern confirmed its higher expression in intercalated nidopallium regions relative to intercalated hyperpallium regions (**Fig. 5f**), the 57^th^ such ranked gene (**Table S2**). We note that in searching through many *in situ* hybridization profiles, it was difficult to find constitutively expressed genes that differed between the two mesopallium regions and between the nidopallium and hyperpallium regions, consistent with the RNA-Seq findings.

### 3.5 Functional gene networks in specific avian telencephalic populations

The results from the pairwise differential expression analysis highlight shared expression profiles in subdivisions, suggesting an organizational continuum around the ventricle **(Fig 4c).** Investigating co-expression networks of these genes could offer insights into whether the subdivisions above and below the ventricle also exhibit a functional continuum, with similar genes performing similar functions in related subdivisions, or if there are functional distinctions between each subdivision regardless of shared expression profiles. To test for this possibility, we performed whole gene co-expression network analysis (WGCNA) treating all samples independently. WGCNA finds patterns of co-expression across all genes in the dataset and defines blocks or modules of genes that fluctuate together, often with functional significance (Langfelder and Horvath, 2008; Oldham, Horvath and Geschwind, 2006). If the distinction hypothesis is correct, we would expect to find distinct gene modules for each of the proposed unique subdivisions. However, the presence of gene modules that significantly correlate with subdivisions above and below the ventricle would be strong evidence for the continuum hypothesis. We constructed our networks with a criterion of a 100-gene module minimum in order to avoid small modules driven by single samples and to obtain the most robust findings. We summarized each module by their eigengenes (average expression of all module genes) and tested the strength of each module correlation to each brain region/subdivision. Finally, we tested the functional enrichments of these modules using Gene Ontology analysis. We noted instability in module membership when the intercalated pallial modules were included, potentially due greater divergence between the two populations, and thus we performed network analyses with and without the intercalated pallium regions included.

Without the intercalated pallium regions, selecting a soft power (6) to maximize mean connectivity between genes (**Fig. 6a**), we found a total of 47 modules (**Fig. 6b**). Among these 47 modules, we found five with highly significant positive correlations (r^2^ > 0.9, q < 0.0001), and all grouped according to brain subdivisions (**Fig. 7a**). They included a mesopallium-specific module of dorsal and ventral regions (module 15; **Fig. 7a**). This mesopallium module consisted of 363 genes (**Fig. 8a**), and was highly specialized for functions in lymph vessel development and anatomical structure development (**Fig. 8f**). They included a nidopallium/hyperpallium-specific module (module 17; **Fig. 7a**), consisting of 335 genes (**Fig. 8b**), with functional enrichments in regulation of development growth and anatomical structure development (**Fig. 8f**). There were two arcopallium-specific modules (module 3 and module 5; **Fig. 7a**), consisting of 1,501 (**Fig 8c**) and 1,205 genes, with distinguishing functional specializations of anatomical structure development and regulation of intracellular signaling, respectively (**Fig. 8f**). Finally, we observed and a striatum-specific module (module 1; **Fig. 7a**), consisting of 2,239 genes (**Fig. 8d**), and with a distinguishing specialization for neurogenesis and nervous system development (**Fig. 8f**).

**FIGURE 6.**
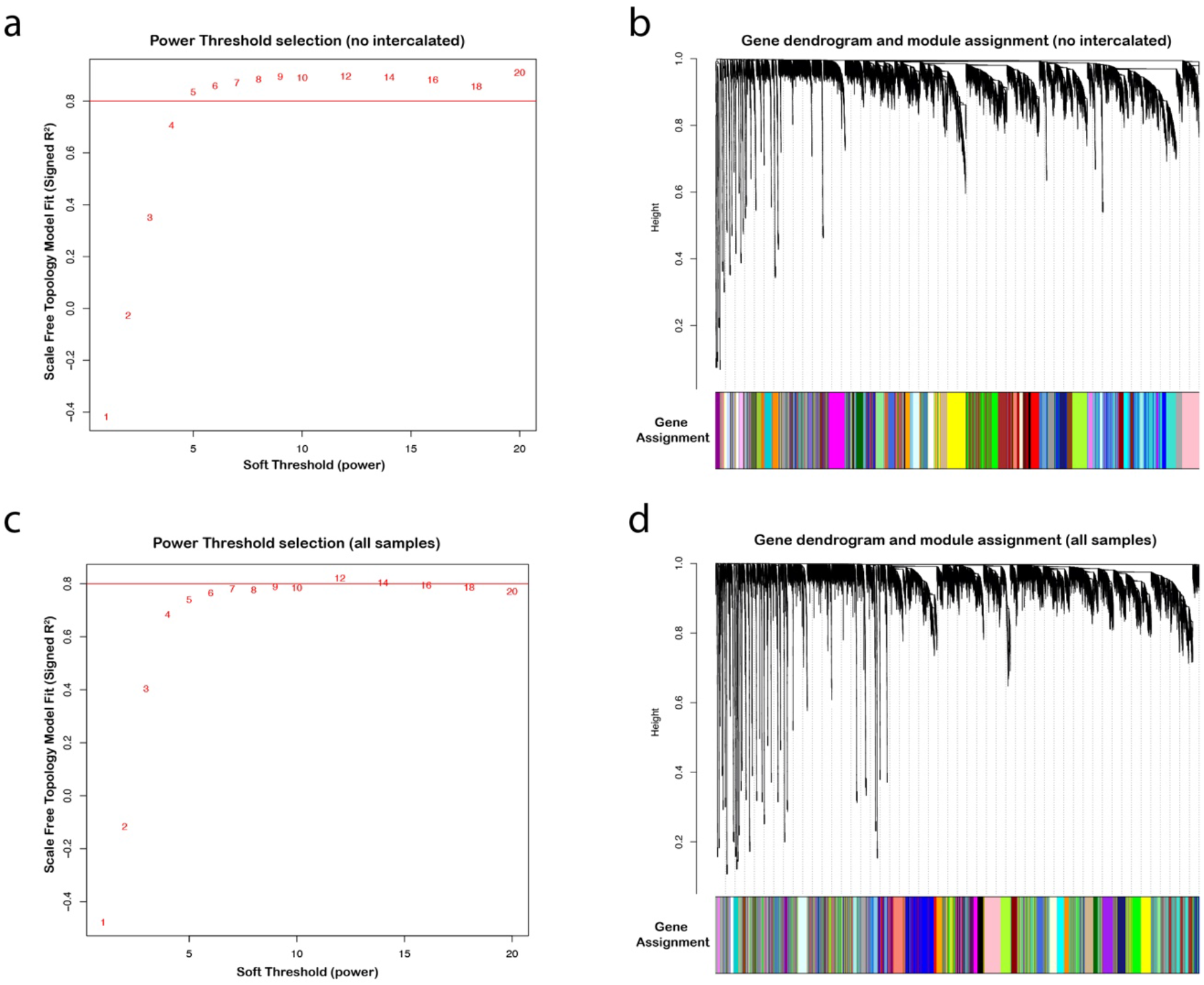
Soft power threshold and gene dendrogram for gene co-expression networks. (**a**) For the network excluding the intercalated regions, a soft power (6) was selected to maximize mean connectivity between genes (at least 80%). (**b**) Gene coexpression network dendrogram drawn from the soft power threshold in (a), resulting in 47 unique modules (colors). (**c**) For the network including the intercalated regions, a soft power (8) was selected to maximize mean connectivity between genes (at least 80%). (**d**) Gene co-expression network dendrogram drawn from the soft power threshold in (**c**), resulting in 38 unique modules.

**FIGURE 7.**
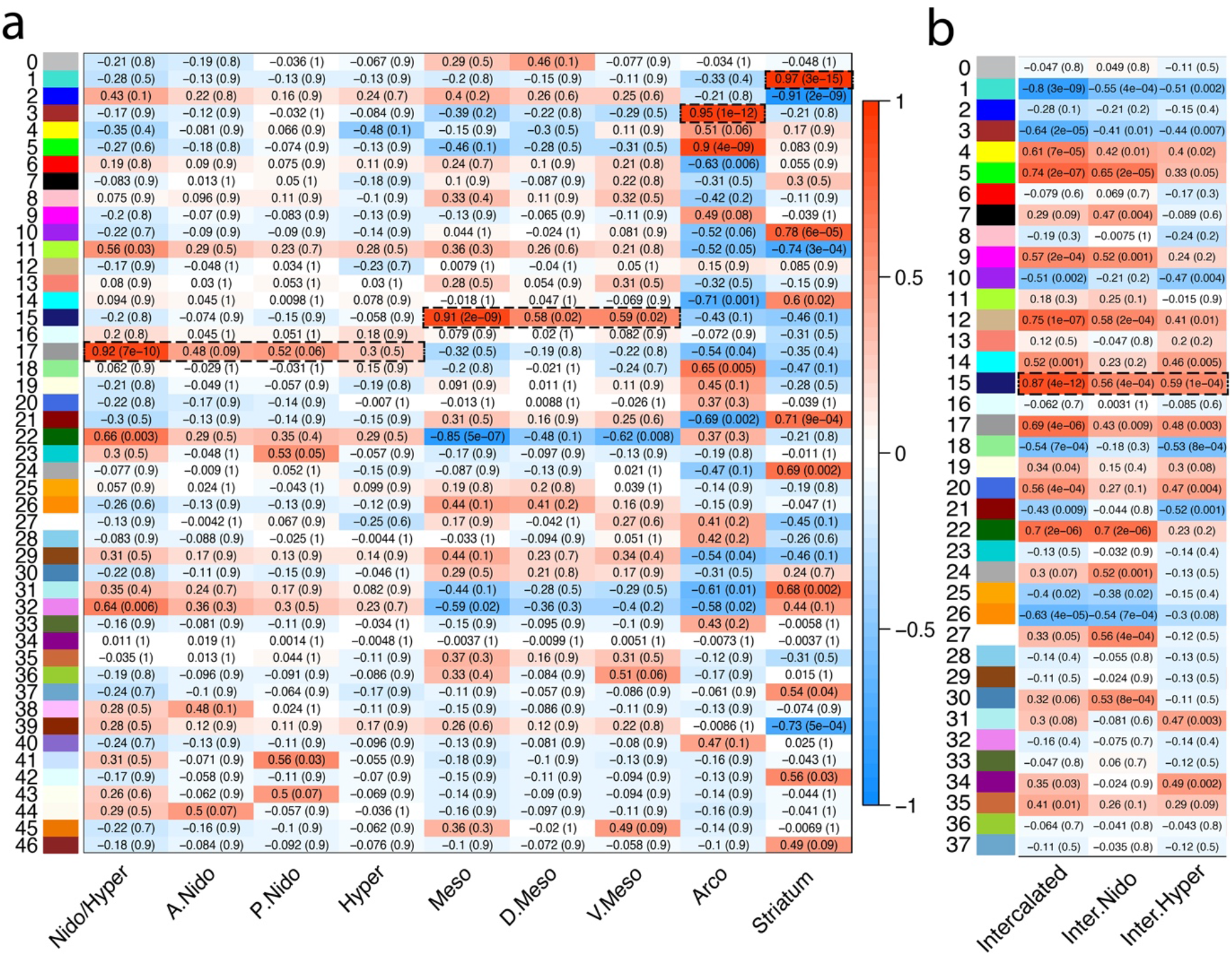
Statistical correlation results of all gene module co-expression networks. (**a**) Module eigengene vectors and subdivision correlations with intercalated pallium regions excluded. The 47 subnetworks in the gene expression data are identified with unique module number and color (left), as in (**b**). Entries show Pearson correlation and associated corrected q-value (parenthesis), testing for statistical relationships between each module eigengene to a unique subdivision or combination of subdivisions. Color scale indicates strength of positive or negative correlation. Dashed boxed regions highlight strong (r^2^ > 0.9) and highly significant (p < 0.0001) correlations. **(b**) Module eigengene vectors and subdivision correlations with intercalated pallium regions included. The 38 subnetworks in the gene expression data are identified with unique module number and color (left), as in Figure 7d; only results for the intercalated regions are shown. Dashed boxed regions highlight strong (r^2^ ~0.9) and highly significant (p < 0.0001) correlations.

**FIGURE 8.**
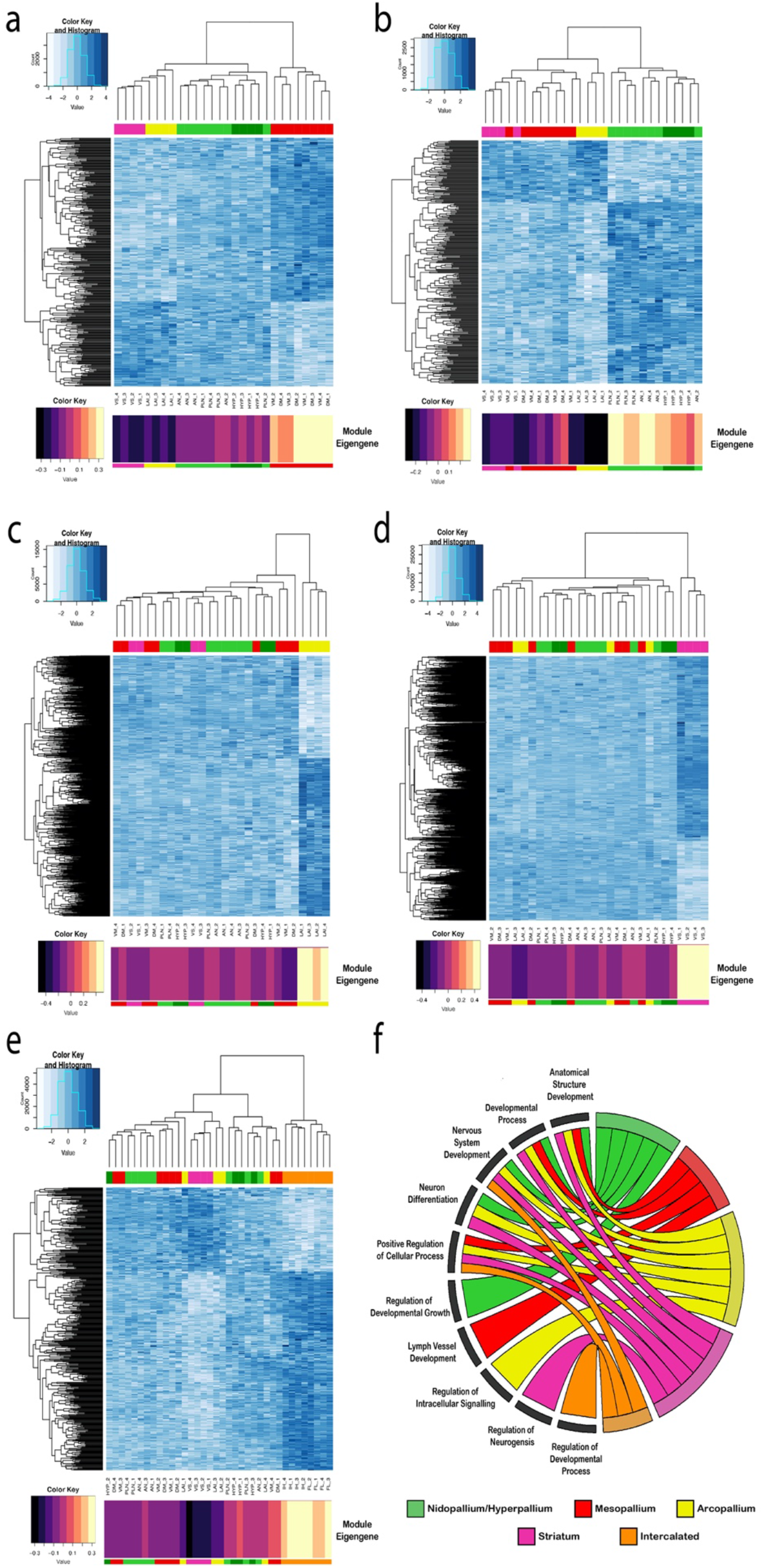
Gene expression module profiles specific to each major avian telencephalic subdivisions and combinations of subdivisions profiled. **(a-e)** Heatmaps of gene expression for subdivision-specific modules. All module specific genes (rows) are plotted; degree of blue indicates level of median scaled gene expression for all module genes. To the left is the gene expression tree dendrogram; at the top is the brain region (colors) dendrogram relationships for each sample (bird). At the bottom is module eigenvector value for each subdivision module; the larger the number (or lighter color), the stronger the relationship of that sample to the module eigengene. (**a**) Mesopallium-specific (MV, MD; red) module (15, nGene = 363). There is a strong anticorrelation between this module’s genes and the arcopallium and striatum. (**b**) Nidopallium (PLN, AN; lightgreen) and hyperpallium (HYP; darkgreen) specific module (17, nGene = 335). (**c**) Arcopallium (LAI) specific module (3, nGene = 1,501). (**d**) Striatum (VS; maroon) specific module (1, nGene = 2,239). (**e**) Intercalated-specific (IH, IN; orange) specific module (15, nGene = 442 (**f**) Chord diagram of significant GO terms for each neural subdivision module. Each subdivision module contains specific functional enrichments (bottom left quadrant), as well as substantial overlap in function for nervous system development and neuron differentiation (top left quadrant). A list of the most significant GO terms can be found in **Table 2**, with a complete list in **Table S3 (a-d)** and **Table S4 (e)**.

When including the intercalated regions in the WGNCA, we obtained fewer modules (38: **Fig. 6c-d**), but there was one additional module with a highly significant correlation (r^2^ = 0.87, q < 0.0001). This intercalated pallium-specific module (module 15; **Fig. 7b**), consisting of ~1,700 genes (**Fig 8e**), had a distinguishing functional specialization for regulation of developmental processes (**Fig. 8f**). Intriguingly, while each module exhibited a functional enrichment specific to each subdivision, there were five development-specific enrichments shared between most or all of the brain subdivisions: anatomical structure development; developmental process; neuron system development; neuron development; and positive regulation of cellular process (**Fig. 8f; Table 2**). Importantly, these similar functional modules were composed of mostly non-overlapping gene sets for each brain subdivision (**Table S3, S4**), suggesting these shared functional specializations are being achieved with unique sets of genes.

**TABLE 2:** Top 5 gene ontology terms and p-values for the subdivision-specific modules. All subdivision-specific modules are enriched for either anatomical structure or nervous system development, but with different genes, suggesting these regions utilize different genes for the same functions to distinguish themselves from each other during development.

| Subdivision | GO Term | P-Value |
| --- | --- | --- |
| Nidopallium/Hyperpallium | neuron development | 1.50e-03 |
|  | nervous system development | 1.75e-03 |
|  | system development | 1.79e-03 |
|  | multicellular organism development | 1.83e-03 |
|  | anatomical structure development | 3.49e-03 |
| Mesopallium | lymph vessel development | 4.27e-05 |
|  | regulation of system process | 8.88e-05 |
|  | multicellular organismal process | 5.31e-04 |
|  | regulation of multicellular organismal process | 8.64e-04 |
|  | lymphangiogenesis | 2.22e-03 |
| Arcopallium | system development | 4.63e-12 |
|  | anatomical structure development | 5.98e-12 |
|  | multicellular organism development | 2.40e-11 |
|  | developmental process | 8.58e-11 |
|  | anatomical structure morphogenesis | 1.15e-10 |
| Striatum | regulation of nervous system development | 8.91e-09 |
|  | developmental process | 1.25e-08 |
|  | generation of neurons | 3.25e-08 |
|  | anatomical structure development | 3.37e-08 |
|  | neurogenesis | 4.71e-08 |
| Intercalated | positive regulation of nervous system development | 6.82e-04 |
|  | positive regulation of cellular process | 2.50e-03 |
|  | positive regulation of biological process | 8.60e-03 |
|  | cellular component organization | 8.95e-03 |
|  | regulation of multicellular organismal development | 9.19e-03 |

Importantly, with or without the intercalated pallium samples included, there were no modules where the dorsal/ventral mesopallium, the nidopallium/hyperpallium, or the intercalated regions exhibited strong separate gene network correlations (at r^2^ ~ 0.9, p < 0.0001; **Fig. 7a,b**). These individual regions had weaker correlations (r^2^ 0.4-0.6, p < 0.05), but mainly in the same module that the complementary regions above and below the ventricle had, and with a stronger correlation when grouped together (**Fig. 7a,b**). The hyperpallium had no significant positive correlation at all (p > 0.1) and one weakly significant negative correlation (r^2^ = 0.48, p = 0.01; meaning absence of this network; **Fig. 7a**), indicating that the hyperpallium cannot be distinguished from the nidopallium in terms of gene functional networks in this analysis. The higher correlations in the combined regions cannot be explained by higher sample numbers alone, because if these regions above and below the ventricle were significantly different, the correlations would be weakened, not strengthened, by combining them. The strengthening demonstrates shared functional molecular properties.

We also noted some intriguing higher order relationships among some brain subdivision-specific modules. The mesopallium-specific module had an inverse gene expression relationship of the same interacting genes in the arcopallium and striatum (negative correlations in module 15 in **Fig. 7a**; expression profile in **Fig. 8a**). A similar finding was seen for the nidopallium/hyperpallium module 17 relative to the arcopallium (**Figs. 7a, 8b**). In contrast for the arcopallium-specific (module 3) and striatum-specific (module 1) modules, expression of the genes in the other brain regions were more uniform (**Figs 7a, 8c,d**). This suggests that there are broad programs of gene regulation that can be turned up in one brain subdivision and turned down in another.

### 3.6 Hub genes reveal potential master regulators of avian subdivision organization

Each module contains genes with co-regulated expression, but some are more connected than others. Genes with the highest connectivity to other genes in the WGNCA are referred to as hub genes and are the most promising candidates for master regulators of genes in a brain subdivision module (Seo *et al*., 2009), and could offer insights into essential genetic players for the similar populations around the ventricle. To determine hub genes for each subdivision, we calculated the strength of connectivity of each gene within a module (module membership, MM) as well as the strength of the genes expression to the subdivision of interest (gene significance, GS). These can be positive or negative values depending on the gene’s expression relative to all other subdivisions, where one gene’s downregulation might be just as important as another gene’s upregulation. Plotting the absolute value of these metrics reveals the relative contribution of each gene to the module-subdivision association.

We found significant associated hub genes specific to each brain subdivision (**Table S5**) and visualized the top 50 in network diagrams (**Fig. 9a-e**; defined as genes in the upper quadrant, with an absolute value of MM > 0.80 and an absolute value of GS > 0.80). Each subdivision had several hub gene transcription factors with various ranges of downstream target genes, which could be strong candidates for regulating the other gene within the module. For example, the mesopallium module exhibited moderate interconnectivity (median n = 7) between its top 50 hub genes. One of the most densely connected genes (individual n = 28) was the *SATB2* transcription factor (**Figs. 5a, 9a**), which is known to be expressed in the superficial layers of the mammalian cortex, and controls the expression of genes involved in intracortical pyramidal neuron connectivity (Cera *et al*., 2019; Alcamo *et al*., 2008). This suggest that this transcription factor may help specialize the dorsal and ventral portions of the mesopallium in a similar manner. The nidopallium/hyperpallium module also exhibited moderate connectivity between its top 50 hub genes (median n = 7), and was similarly defined by strong connectivity (individual n = 16) of *DACT2* (**Fig. 9b**), which regulates intracellular signaling during development (Schubert *et al*., 2014). Two other hub genes in this module were the axon guidance genes *SEMA6A* (individual n = 6) and *EPHA8* (individual n = 10), suggesting together they may drive the shared connectivity motifs (Jarvis *et al*., 2013; Stacho *et al*., 2020). The intercalated pallium (median n = 6.5) had *GRIN2A* ionotropic glutamate receptor and the doublecortin kinase *DCLK1* specific to the intercalated nidopallium **(Fig. 9c**), suggesting these genes may control the specialized signaling pathways for this region. The arcopallium module exhibited many more connections between hub genes (median n = 24). Two of the most interconnected genes in the arcopallium (**Fig. 9d**) are well-known transcription factors *LHX9* (individual n = 25) and *ETV1* (aka *ER81*, individual n = 39) previously studied in the avian brain (Jarvis *et al*., 2013), and expressed in deep layer projection neurons and the pallial amygdala of mammals (Dugas-Ford, Rowell and Ragsdale, 2012; Abellán, Desfilis and Medina, 2013), indicating that the transcription factors that define the intermediate arcopallium may have been long-discovered. Likewise, the top hub genes for the striatum exhibited strong interconnectivity (median n = 25), and included well-known dopamine receptors (*D1A, D1B*; aka *DRD1* and *DRD5*, respectively), and the *FOXP2* transcription factors (in the top 75 hub genes) that define the striatum (**Fig. 9e**) (Haesler *et al*., 2007; Teramitsu *et al*., 2010; Kubikova *et al*., 2014). Importantly, while uncharacterized genes (LOC IDs) were replaced with functional aliases whenever possible (see *Methods*), each subdivision-specific hub network was composed of some genes of unknown function, many of which are ncRNAs (22-82% in top 50 hubs), highlighting further need for investigations into the roles of these genes in the differentiation of neural subdivisions. Overall, these analyses demonstrate that the molecular functions of the regions above the ventricle are informative for those below, and vice versa.

**FIGURE 9:**
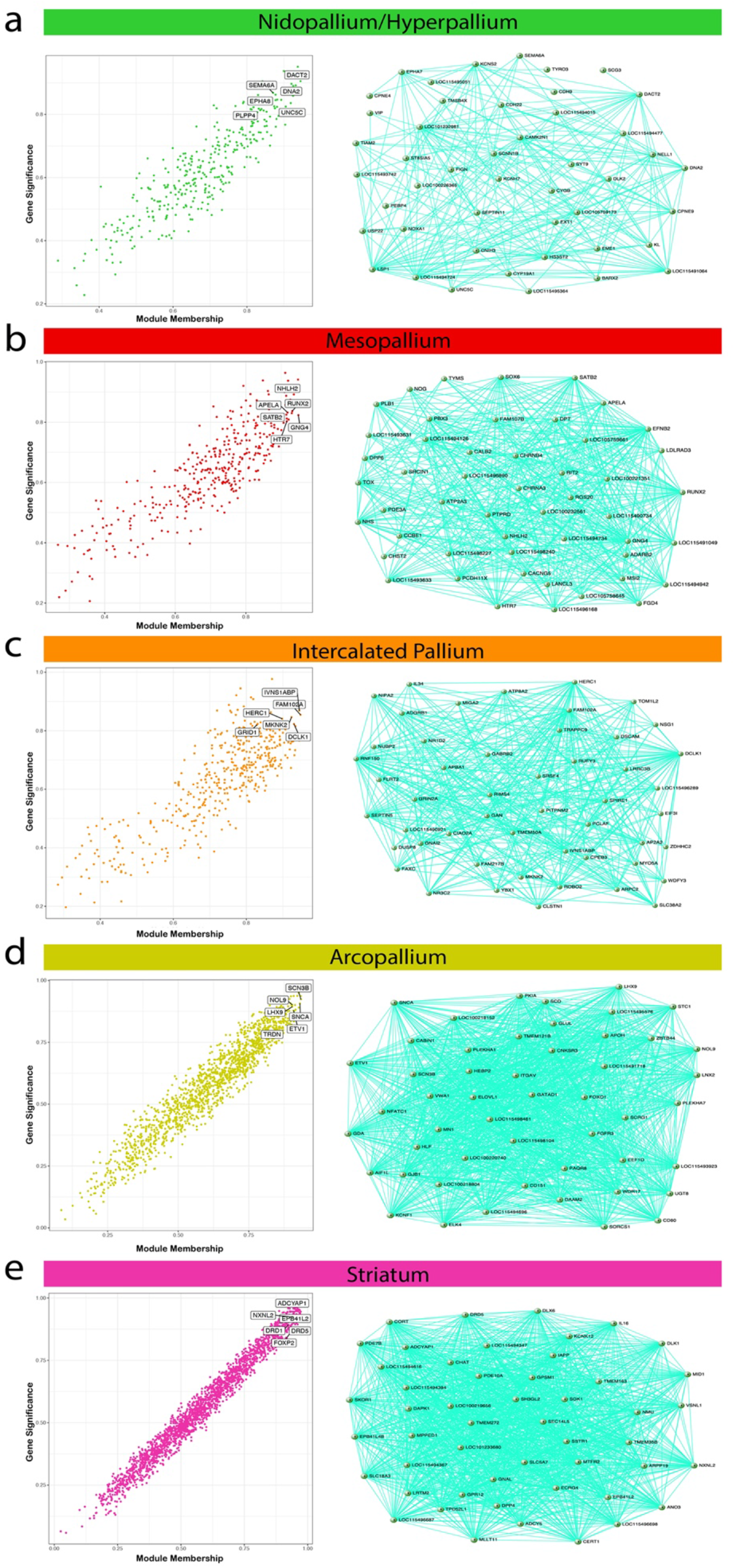
Hub gene identification and top 50 hub gene networks reveal best candidates for subdivision specific expression regulation. (**a-e**) Correlation between module membership (MM) and gene significance (GS) for the (**a**) dorsal/ventral mesopallium, (**b**) nidopallium/hyperpallium, (**c**) intercalated pallium, (**d**) arcopallium, and **(e)** striatum modules. Highlighted are example hub genes with MM > 0.80 and GS > 0.80. The top 50 hub genes and their connections are visualized adjacent to each subdivision-specific module correlation plot. The genes with the highest connectivity offer prime candidates for key regulators of module expression for each subdivision. A full list of hub genes for each subdivision-specific module can be found in **Table S5**.

## 3. DISCUSSION

By examining the entire annotated transcriptome of the major cell populations of the adult avian pallium and striatum at baseline, we demonstrate that the distinction hypothesis, which states that the pallium above the ventricle is a ‘hyperpallium cluster’ distinct in character and cell types from the so-called DVR below the ventricle (**Fig. 1a**), is no longer tenable. Rather, for each major population above the ventricle divide there is a corresponding population below it. The mesopallium regions are the most similar, followed by the hyperpallium and nidopallium, while the intercalated pallial regions are the most divergent. The levels of similarity correlate with the brain region’s relative proximity to the ventricle divide. The findings provide strong support for the continuum hypothesis of avian dorsal and ventral pallium organization (**Fig. 1b**), in which six previous distinctly-named cell populations are really three continuous cell populations that wrap around the ventricle (Chen *et al*., 2013; Jarvis *et al*., 2013).

We believe our findings and interpretations differ from some past individual gene expression and broader transcriptome studies (Belgard *et al*., 2013; Montiel and Molnár, 2013; Montiel *et al*., 2016; Watson and Puelles, 2017; Puelles *et al*., 2016) for several key reasons. First, we had a set of *in situ* hybridization gene expression profiles, Nissl staining, and myelin staining (Jarvis *et al*., 2013; Chen *et al*., 2013; Karten *et al*., 2013) that helped guide understanding of brain population boundaries, from which to perform our dissections for RNA-Seq analyses. This reduced the possibility of contaminating brain region populations in our dissections. Second, we performed laser capture microdissections on thin 12 ∝m sections, as opposed to more gross dissections on thick sections, to reduce cross contamination between brain subdivisions. Third, we kept animals in quiet control conditions to bring brain gene expression to a baseline and consistent state across animals, as opposed to freely behaving animals, where up to 10% of the genes in the genome can be regulated across different cell types within a forebrain circuit (Whitney *et al*., 2014). Even with this control of animal state, we still found cases of many co-regulated genes across brain regions specific to one or two animals, which strongly impacted results when not corrected for. We believe these careful controls lend greater confidence to the results of our experiments.

A case study of these considerations is on the *NR4A2* transcription factor. Puelles et al. (2016) used this gene’s expression pattern to argue that the continuum hypothesis was not plausible, because it labelled only mesopallium below the ventricle, and not above. However, fate mapping of cells in the mesopallium did not support their hypothesis (Bruguier *et al*., 2020) and our companion study (Biegler et al., submitted) found that *NR4A2* is an activity-dependent gene, whose expression changes in different brain regions according to different behaviors, including in both ventral and dorsal mesopallium. In the present study, we found that *NR4A2* is a hub gene in the arcopallium-specific module with no detectable expression in the mesopallium at baseline, highlighting the importance of utilizing animals with well-controlled behavior states for interpretations of gene markers of cell types. Future studies on finer delineations within each subdivision, such as within the arcopallium (Mello *et al*., 2019), would help determine which cell types utilize this gene at rest or during behavior.

The shared molecular gene functions of the brain populations above and below the ventricle contribute to a broader understanding of avian brain subdivision functions. Since all the subdivision-specific modules contain non-overlapping gene sets involved in anatomical structure and nervous system development, these genes are candidates to control the principle neural connectivity and structure differences between subdivisions. For example, the intercalated pallium hub genes are prime candidates involved in forming and maintaining the specialized connections with sensory input neurons from the thalamus (Wang, Brzozowska-Prechtl and Karten, 2010; Jarvis *et al*., 2013). The nidopallium/hyperpallium hub genes specialized in regulation of developmental growth are candidates for growth and maintenance of the cell types in this region. This could include the axon guidance cues like *SEMA6A* and *EPHA8*, perhaps giving rise to the shared connectivity patterns observed in these regions (Jarvis *et al*., 2013). The mesopallium genes enriched in lymph vessel development was a surprise for us and indicates that perhaps there is a relationship between lymph vessel cell types and higher-level brain functions not previously appreciated. The arcopallium genes enriched in intracellular signaling are prime candidates for maintaining the high firing rates and subsequent signaling pathways of these neurons. The striatum hub-specialized specialized for neurogenesis are prime candidates for the high levels of neurogenesis seen in the avian striatum (Paredes *et al*., 2016). Future investigations on single cell transcriptomes within each brain subdivision will be able to determine which cell subpopulations are specialized for these specific sub-functions.

Briscoe et al. (2018) performed RNA-Seq on chicken brains on a subset of brain regions we profiled here in the zebra finch, and also found that the dorsal and ventral mesopallium are similar to each other, and further that each contained cell types similar to the intratelencephalic neurons (IT) of the mammalian cortex. They noted that the transcription factor *SATB2* was a marker for these IT cells in mammals and birds, and is likely an important regulator of genes in this cell type. Our findings suggest that *SATB2* is a centralized hub gene in the mesopallium-specific module that defines both dorsal and ventral regions, suggesting the principle cell type driving the signal in our bulk data are these IT-like cells. Putting these results together, it suggests that our subdivision-specific modules are driven by common cell types. Furthermore, we noted cellular diversity within an established subdivision not seen in its mirror image counterpart, the sparse *SATB2+* cell population in the hyperpallium. Future studies utilizing single cell/nuclei sequencing (Aevermann *et al*., 2018) will further our understanding of cell types within each population surrounding the ventricle, and their evolutionary relationships to cell types in the mammalian brain.

What is the nature of the relationship of these continuous cell populations? Montiel and Molnar (2013) suggest that the similarities in 50 genes seen between the regions above and below the ventricle in the Jarvis et al and Chen et al 2013 studies could be due to convergent evolution of cell types within these cell populations. One rationale for a convergent organization is that nonavian reptile brains have a DVR thought to be homologous to birds whereas their dorsal pallium above the ventricle, sometimes called the ‘dorsal cortex’, is much thinner, and more layered than it is in birds, and thought are more specific to non-avian reptiles (Dugas-Ford, Rowell and Ragsdale, 2012). According to an independent evolution hypothesis, birds would have inherited this layered dorsal cortex organization from their reptile ancestor and then convergently evolved the same cell types as below the ventricle, with pallial thickening similar to the DVR. Under this view, one could imagine, that a proportion of genes and cell types evolved convergently between the Wults and DVR. However, in this study, by examining over 20,000 genes, we found that the hyperpallium and nidopallium differ by only 35-50 genes (~0.2%) depending on the portion sampled, and the mesopallium regions surrounding the ventricle differ by only 4 genes (0.02%). Such dramatic similarity in expression between these regions seems more parsimonious with shared homology than large-scale, transcriptome-wide convergence. Furthermore, the presence of shared modules between the dorsal and ventral portions of the mesopallium, between the nidopallium and hyperpallium, and between the intercalated regions, suggest that these brain subdivisions share large-scale organization in gene expression networks. Such network architecture is difficult to evolve convergently, and is more often taken as evidence of shared functional networks in homologous brain regions (Oldham, Horvath and Geschwind, 2006). Investigating the open regulatory regions in the continuous cell populations using ATAC-Seq would offer insights into the regulatory landscape of the cell types within these populations and if they are activating the same expression networks via convergent or homologous means. Systematic lineage tracing experiments, paired with *in situs* for the shared expression markers found in this study, would provide evidence to test competing hypotheses on the developmental origins of similar subdivisions (Solek and Ekker, 2012; Montiel *et al*., 2016; van Essen *et al*., 2020).

Not only do our findings provide a deeper understanding of the avian brain organization, they have important implications for understanding the homology and evolution of pallial cell populations in birds relative to other non-reptile vertebrates. At a minimum, evolution-based hypotheses using gene expression profiling should consider the pallial populations above and below the ventricle together. The intercalated nidopallium has been proposed to be homologous to layer 4 thalamic recipient neurons of the mammalian cortex, based on gene expression profiling (Dugas-Ford, Rowell and Ragsdale, 2012; Jarvis *et al*., 2013); our findings indicate that this would then also apply to the intercalated hyperpallium of birds. The nidopallium and ventral mesopallium populations have been proposed to be homologous to cell types in the upper layers 2 and/or 3 of the mammalian cortex, respectively (Wang, Brzozowska-Prechtl and Karten, 2010); if so, then the hyperpallium and dorsal mesopallium populations of birds may also be considered homologous to cell types in these same layers of the mammal cortex. The same logic applies to the hypotheses that propose that the avian pallial regions ventral to the ventricle are homologous to the mammalian claustrum and amygdala (**Fig. 1a**). If true, then it will be difficult to justify the claim that the avian dorsal regions are in turn homologous to the 6-layered cortex separate from the claustrum and amygdala.

The shared transcriptomes and molecular functions found in this study, combined with the shared neural connectivity motifs and developmental origins found in previous studies (Jarvis *et al*., 2013; Chen *et al*., 2013), provide a strong rationale for the mesopallium and intercalated pallium regions above and below the ventricle to have the same population names. Furthermore, our results indicate that the hyperpallium and nidopallium should also have the same population name. However, from a practical perspective, more revised renaming using similar names, such as dorsal and ventral nidopallium or dorsal and ventral hyperpallium, could cause more confusion in the literature. One alternative approach is to apply a new alternative naming system, such as that proposed in the Jarvis and Chen 2013 studies, based on known differences in neural connectivity among the avian pallium subdivisions. Here the intercalated hyperpallium and intercalated nidopallium (L2, entopallium, and basorostralis) would be called the dorsal and ventral primary (1°) pallium, respectively, since they are the first recipient population of thalamic sensory input into the telencephalon (**Fig. 1b**). The hyperpallium and nidopallium would be called the dorsal and ventral secondary (2°) pallium, respectively, as they receive their main extra-telencephalic input via of the 1° pallium regions. The dorsal and ventral mesopallium would be called the dorsal and ventral tertiary (3°) pallium, respectively, as they receive their main extra-telencephalic input via the 2° pallium. The arcopallium would be called the quaternary (4°) pallium, as it contains cell populations that are the main output of the telencephalon.

In conclusion, the highly similar molecular makeup of the populations above and below the ventricle necessitate shared functions, helping to inform our understanding of avian brain organization and allowing for new interpretation and translation of findings between brain subdivisions and between species.

## Acknowledgements

We thank Samara Brown for editing and constructive feedback on the manuscript, and Thomas Carrol of the Rockefeller University Bioinformatics Core Facility for his advice on several analyses. This project was funded by the Howard Hughes Medical Institute (OSU1013377) and Rockefeller University start-up funds to E.D.J., and National Science Graduate Research Fellowship 2015202850 to G.G.

## AUTHOR CONTRIBUTIONS

G.G. performed experiments and analyses and co-wrote the paper. BH generated the RNA-Seq libraries. GD helped with LCM dissections and processing primary pallium regions. MB provided *in situ* hybridization analysis. OF consulted on experimental design and supervised RNA sequencing. EDJ supervised the study and co-wrote the paper.

## DATA AVAILABILITY

All RNA sequencing data supporting the findings of this study are deposited in the NCBI SRA archives upon publication with an accession number listed here.

